# Gibson Assembly-based direct cloning of plasmid DNA in *Lactiplantibacillus plantarum* WCSF1

**DOI:** 10.1101/2022.09.09.507304

**Authors:** Marc Blanch Asensio, Sourik Dey, Shrikrishnan Sankaran

## Abstract

Lactobacilli are gram-positive bacteria that are growing in importance for the healthcare industry and genetically engineering them as living therapeutics is highly sought after. However, progress in this field is hindered since most strains are difficult to genetically manipulate, partly due since their complex and thick cell walls limit our capability to transform them with exogenous DNA. To overcome this, large amounts of DNA (>1 μg) are normally required to successfully transform these bacteria. An intermediate host, like *E. coli*, is often used to amplify recombinant DNA to such amounts although this approach poses unwanted drawbacks such as increase in plasmid size, different methylation patterns and limitation of introducing only genes compatible with the intermediate host. In this work, we have developed a direct cloning method based on Gibson assembly and PCR to amplify the DNA to sufficient quantities for successful transformation in *L. plantarum* WCFS1. This advantage of this method is demonstrated in terms of shorter experimental duration and the possibility to introduce a gene incompatible with *E. coli* into *L. plantarum* WCFS1.

## INTRODUCTION

Lactobacilli are a group of Gram-positive bacteria of great importance to the food and healthcare industries with numerous strains identified as being beneficial for humans, and used as probiotics [1,2,3]. Furthermore, since they naturally colonize almost every site of the human body that hosts a healthy microbiome, *e.g*. the gastrointestinal tract [4,5], urogenital tracts [6], oral cavity [7] and nasal cavity [8], lactobacilli are an excellent foundational candidate for the development of live biotherapeutic products (LBPs) [9]. Beyond their natural health benefits, there is considerable interest in engineering them with heterologous genes for therapeutic applications like drug delivery [10,11] and mucosal vaccinations [12,13]. However, one of the crucial factors slowing down progress in lactobacilli engineering is difficulties in transforming them with exogenous DNA [14]. This is largely due to their thick and complex cell wall structures that can be overcome by using large quantities of plasmid DNA (>1 μg) for transformation [15]. To obtain such high plasmid DNA quantities, shuttle vectors are often used that can be amplified in intermediate hosts, predominantly *E. coli* [16]. To facilitate the construction of recombinant plasmids, several shuttle vectors have been identified which can undergo stable replication in both the cloning host, *E. coli* and the relevant Lactobacilli strains [17,18,19]. Nevertheless, since *E. coli* is a Gram-negative bacterium that is phylogenetically distant from Lactobacillus genera, this strategy can lead to genetic sequence incompatibilities due to GC-content differences [20], DNA methylation [21], repetitive sequence insertions [22] and toxic protein buildup in the *E. coli* cloning host [23]. Alternatively, the Gram-positive lactic acid bacterium, *Lactococcus lactis*, can also be used as an intermediate host for recombinant plasmid construction [24]. However, the availability of functional replication origins in *L. lactis* is limited [25] and inclusion of additional broad-range replicons significantly increases the size of the plasmid. The excessive increase in the size of the plasmid might lead to segregational instability [26] and thereby limits the size of the heterologous genes that can be included in it. Hence, it is desirable to be able to clone these lactobacilli without the need for an intermediate host. To avoid the need for an intermediate host, Spath et al. developed a direct cloning approach based on the assembly of PCR-amplified DNA fragments by restriction digestion and ligation to obtain sufficient quantities for transformation [27]. While this approach was very successful in transforming *Lactiplantibacillus plantarum* CD033, the method requires the presence of restriction sites within the DNA sequences, which can limit the versatility of combining heterologous genes in the plasmid.

In this work, we report a direct cloning method that leverages the Gibson assembly strategy and takes advantage of recent advances in cost-effective oligonucleotide synthesis. By doing so, we avoid the need for restriction sites and improve the feasibility of combining diverse DNA sequences to construct versatile recombinant plasmids. We demonstrate this direct cloning method in *Lactiplantibacillus plantarum* WCFS1, one of the most commonly engineered probiotic *Lactobacillus* strains for which improved engineering methods are highly sought [28,29]. Furthermore, this direct cloning method is considerably quicker in comparison to indirect cloning methods requiring an intermediate host. We have characterized the efficiency and accuracy of this approach and have demonstrated the successful cloning of a genetic segment expressing the medically relevant protein, Elafin which showed a high failure rate when being cloned through the intermediate host, *E.coli*. Thus, this direct cloning method will be instrumental in enabling the cloning of Lactobacilli with a wider variety of heterologous genes and with greater versatility than previously possible.

## MATERIALS AND METHODS

### Bacterial strains and growth conditions

*L. plantarum* WCFS1 was used as the parent strain in this study. The strain was grown in the De Man, Rogosa and Sharpe (MRS) media (Carl Roth GmbH, Germany, Art. No. X924.1). Recombinant *L. plantarum* WCFS1 strains were grown in MRS media supplemented with 10 μg/mL of erythromycin (Carl Roth GmbH, Art. No. 4166.2) at 37 °C and 250 rpm shaking for 16 h. For the indirect cloning experiments, NEB 5-alpha Competent *E. coli* cells were used (New England Biolabs GmbH, Germany,Art. No. C2987). This strain was grown in Luria-Bertani (LB) medium (Carl Roth GmbH, Art. No. X968.1). Recombinant *E. coli* DH5α strains were grown in LB media supplemented with 200 μg/mL of erythromycin at 37 °C,250 rpm shaking conditions for 16 h.

### Molecular Biology

Q5 High Fidelity 2X Master Mix (New England Biolabs GmbH [NEB], Germany, No. M0492S) was used to perform DNA amplification. Amplified DNA products were purified using the Wizard^®^ SV Gel and PCR Clean-Up System (Promega GmbH, Germany, Art. No. A9282). In all agarose gels, the 1 kb Plus Ladder from

ThermoFisher was run as a reference standard (No. 10787018). Primers were synthesized by Integrated DNA Technologies (IDT) (Louvain, Belgium) and the elafin gene fragment was ordered as eBlock from IDT (Coralville, USA). All primers used in this work are shown in Supporting Information Figure S1. The mCherry gene fragment was amplified by PCR from a plasmid previously generated in our lab. The genetic sequences of mCherry and elafin genes are shown in Supporting Information Figure S2. The plasmid *pLp3050sNuc* (Addgene plasmid # 122030) [30] was used as the vector backbone for the recombinant plasmids generated in this study. The Codon Optimization tool from IDT (Choice Host Organism – *L. acidophilus*) was used to optimize the codon bias for mCherry coding segment. The Java Codon Adaptation Tool (JCat) [31] was used to codon-optimize the gene encoding for the human peptidase inhibitor 3, elafin (GenBank ID: AAX36874.1) using the codon optimization database for *L. plantarum* WCFS1. DNA assembly was performed using the HiFi Assembly Master Mix (NEB GmbH, Germany, Art. No. E5520S). For plasmid circularization, the Quick Blunting Kit (NEB GmbH, Germany, Art. No. E1201S) and the T4 DNA Ligase enzyme (NEB GmbH, Germany, Art. No. M0318S) were used.

### *L. plantarum* WCFS1 Electrocompetent Cell Preparation

Wild-type *L. plantarum* WCFS1 was cultured overnight in 5 mL of MRS media and at 37 °C with shaking (250 rpm). After 16h, 1 mL of the culture (OD_600_ = 2) was added to 20 mL of MRS media and 5 mL of 1% (w/v) glycine. This secondary culture was incubated for roughly 4 h at 37 °C and 250 rpm until OD_600_ reached 0.8. The cells were then harvested by centrifugation at 4000 rpm (3363 X g) for 12 min at 4°C. After manually discarding the supernatant, the pellet was washed twice with 5 mL of ice-cold 10 mM MgCl2. After that, the pellet was washed twice with ice-cold Suc/Gly solution (1 M sucrose and 10% (v/v) glycerol mixed in a 1:1 (v/v) ratio), first with 5 mL and second with 1 mL. Next, the supernatant was manually discarded, and the bacterial pellet was resuspended in 450 μL of ice-cold Sac/Gly solution. Finally, 60 uL aliquots were prepared and immediately used for DNA electroporation or stored at −80 °C for future use.

### Electroporation based Transformation in *L. plantarum* WCFS1

For electroporation transformation, plasmids were first mixed with 60 μl of electrocompetent cells at quantities (300 – 1200 ng) specified in the Results section. After a short incubation on ice, the mixture was transferred to ice-cold electroporation cuvettes with a 2 mm gap (Bio-Rad Laboratories GmbH, Germany, #1652086). Electroporation was performed using the MicroPulser Electroporator (Bio-Rad Laboratories GmbH, Germany), with a single pulse (5 ms) at 1.8 kV. Immediately after the pulse, 1 mL of room-temperature MRS media was added, and the mixture was pipetted into a 1.5 Eppendorf tube and incubated at 37 °C and 250 rpm for 3 h. Following this incubation, cells were pelleted down at 4000 rpm (3363 X g) for 5 min. 800 μL of the supernatant was discarded, and the remaining 200 μL was used for cell resuspension by slow pipetting. Finally, the resuspended pellet was plated on MRS Agar plates supplemented with 10 μg/mL of Erythromycin, and plates were incubated at 37 °C for 48 h for colonies to grow.

### Direct cloning method in *L. plantarum* WCFS1

This study created and optimized a novel direct cloning method based on amplifying and circularizing Gibson-assembled gene fragments to obtain adequate quantities of plasmid DNA that were then directly transformed in *L. plantarum* WCFS1 (Figure 1). For the HiFi Assembly reaction, complementary overhangs were included by PCR using a set of primers that contained the corresponding overhangs at the 5’ ends. In the HiFi assembly reaction, 50 ng of the PCR-amplified linear vector with overlapping DNA fragments and 10 ng of the corresponding eBlock were mixed along with 10 μl of the HiFi DNA Assembly Master Mix (Mili-Q water added up to 20 μl). The reaction was incubated at 50 °C for 30 minutes. After that, 5 μL of the assembled product was used as a template for an additional PCR, using a set of primers that annealed to the insert (eBlock). The final volume of this PCR was 120 μl, and the amplification cycle threshold was set at 22. 5 μl of this reaction was run on an agarose gel to confirm amplification (Supporting Information Figure S3). After purifying the linear PCR product, 3500 ng of DNA were phosphorylated using the Quick Blunting Kit. This reaction was performed as suggested in the standard reaction protocol. 2.5 μl of the 10X Quick Blunting buffer and 1 μl of the Enzyme Mix were mixed with the purified DNA (3500 ng). Milli-Q water was added up to 25 μl. The reaction was incubated for 30 minutes at 25 °C to allow the reaction to occur and then for 10 minutes at 70 °C to inactivate the enzymes. Next, phosphorylated DNA was ligated using the T4 ligase enzyme. This reaction was slightly modified from the standard protocol because a higher amount of DNA was added to the reaction. Per ligation, 500 ng of phosphorylated DNA (3.6 μl of the Quick Blunting reaction) were mixed with 1.5 μl of T4 Ligase enzyme and 2.5 μl of 10X T4 Ligase Buffer. Milli-Q water was added up to 25 μl. The number of ligation reactions depended on the amount of DNA intended to be circularized. The ligation reactions were incubated for 2.5 hours at 25 °C and then 10 minutes at 70 °C to inactivate the enzymes. After the incubations, ligations were mixed, and the ligated plasmid DNA was purified using the Promega kit. In this purification, DNA was eluted 3 times with 9 μl of Milli-Q water each time to obtain the highest DNA concentration. The DNA concentration of the ligated mixture was measured (absorbance at 260 nm) using a NanoDrop Microvolume UV-Vis Spectrophotometer (ThermoFisher Scientific GmbH, Germany). The purified ligated products were transformed into *L. plantarum* WCFS1 electrocompetent cells.

**Figure 1.**
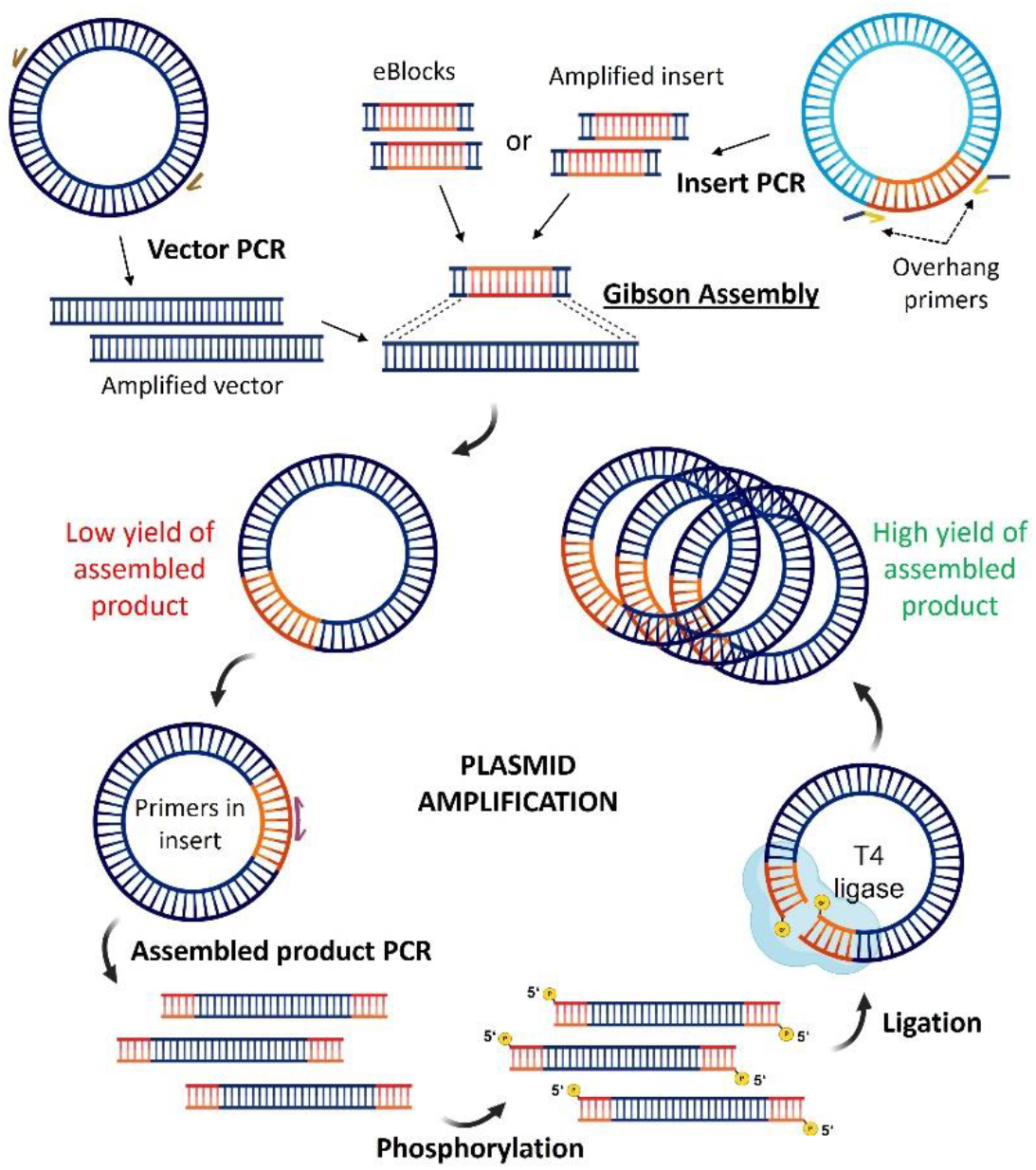
Scheme of the PCR-based plasmid amplification in the direct cloning method. The scheme was generated using BioRender.

Sequence verification was performed by amplifying the target gene segment using PCR to amplify a desired section of our transformed plasmid directly from the bacterial pellet. To do so, the bacteria of interest were inoculated in MRS media supplemented with 10 μg/mL of erythromycin, and the bacteria were incubated overnight at 37 °C and 250 rpm. The following day, 1 mL of the bacterial culture was collected in a 1.5 mL Eppendorf and centrifuged at 4°C for 3 min at 8400 X g. The supernatant was manually discarded, and the residual pellet fraction was scratched off with a sterile pipette tip and used as a template for the PCR (100 μl as final reaction volume, 28 cycles). In the PCR, 10 minutes at 98 °C were set as initial denaturalization. After the PCR, 5 μl of the PCR products were assessed by agarose gel electrophoresis to confirm amplification at the expected size. Next, the PCR product was purified, and the DNA concentration was measured using the Nanodrop. Finally, 2000 ng of the purified PCR product was sent for sequencing to Eurofins Genomics GmbH (Ebersberg, Germany). An additional DNA purification step before Sanger sequencing was employed (Additional Service: PCR Purification).

### Indirect cloning via the intermediate host strain *E. coli*

For the indirect cloning, an additional HiFi reaction was performed (identical to the HiFi reaction set for the direct protocol). However, an additional step was done before setting this reaction, which involved the restriction enzyme digestion with DpnI (NEB GmbH, Germany, Art. No. R0176S). In this reaction, 500 ng of the purified PCR product were mixed with 1 μl of DpnI enzyme and 1 μl of rCutSmart buffer (Milli-Q water was added up to 10 μl). Incubation was performed for 30 minutes at 37 °C followed by 10 minutes at 70 °C. The digested product was used for the HiFi Assembly reaction. Once the HiFi Assembly reaction was done, it was transformed into NEB 5-alpha Competent cells (50 μl). In this transformation, NEB 5-alpha Competent cells were first thawed on ice for 10 minutes. After that, 8 μl of the reaction were properly mixed with the competent cells by pipetting and incubated on ice for 20 minutes Following the incubation, a 60-second heat shock was performed by placing the cells at a 42°C water bath. Next, cells were again incubated on ice for 5 m ice for 5 minutes. After that, 950 μl of SOC media was added, and cells were incubated for 1 hour at 37 C. Finally, 150 μl of the culture was plated in an LB agar plate supplemented with 200 μg/mL of erythromycin and incubated at 37 °C overnight.

For the pLp_mCherry cloning, the screening of positive clones was done using the Gel Documentation System Fluorchem Q (Alpha Innotech Biozym Gmbh, Germany). Bacterial colonies expressing mCherry were imaged in the Cy3 channel (Exλ/Emλ = 554 nm/568 nm) and the corresponding brightfield image was taken using the ethidium bromide channel (Exλ/Emλ = 300 nm/600 nm). One red colony was inoculated in LB media supplemented with 200 μg/mL of erythromycin and incubated at 37 °C overnight. The following day, plasmid extraction was performed using the Plasmid extraction miniprep kit (Qiagen GmbH Germany, Art. No. 27104). The plasmid DNA concentration was measured using the 260 nm absorbance setting on the NanoDrop Microvolume UV-Vis Spectrophotometer. The plasmid was then transformed into *L. plantarum* WCFS1 electrocompetent cells.

For the pLp_elafin cloning, 20 colonies were streaked in a fresh LB agar plate supplemented with 200 μg/mL of erythromycin and incubated at 37 °C overnight. The following day, positive clones were screened by PCR using a forward primer that annealed to the vector and a reverse primer that annealed to the elafin gene (100 μl as final reaction volume, 28 cycles). In the PCR, 10 minutes at 98 °C were set as initial denaturalization. Three positive clones were inoculated in LB media supplemented with 200 μg/mL of erythromycin and incubated at 37 °C overnight. The following day, the respective plasmids were extracted and sent for sequencing to Eurofins Genomics GmbH (Ebersberg, Germany),

## RESULTS AND DISCUSSION

The Gibson assembly approach to combining DNA fragments requires the fragments to contain terminal overhangs that complementarily overlap by ~20 bases. The method was originally developed to stitch together the first artificial genome due to the high level of flexibility it provided compared to restriction digestion-based methods [32]. It is a single reaction assembly method that can be performed without thermal cycling and within an hour. In this work, we employ Gibson-assembly to assemble plasmid constructs for direct cloning in *L. plantarum* WCFS1. As shown in Figure 1, our method involves PCR amplification of a vector and an insert with overlapping arms, followed by their Gibson assembly that yields a low quantity (50-80 ng) of the assembled plasmid. To obtain sufficient quantities for transformation in Lactobacilli, a second amplification and recircularization was performed, yielding >1 μg of the desired construct. This method was characterized and optimized in terms of transformation efficiency, accuracy, and capability for cloning challenging genes in *Lactiplantibacillus plantarum* WCFS1. Different plasmid constructs were assembled and compared with indirect cloning using *E. coli* as an intermediate host.

### Transformation efficiency and accuracy of the direct cloning method

Transformation efficiency (TE) indicates the extent to which cells can take up DNA from the extracellular space and express the genes encoded by it [33]. While it is possible to transform *L. plantarum* WCFS1 with extracellular DNA, TE is typically poor [27]. To demonstrate this, a simple plasmid construct, pLp_mCherry, with gene sequences ideal for indirect cloning through *E. coli* was used. This plasmid consisted of a p256 replicon, an erythromycin resistance cassette and the mCherry gene driven by a strong constitutive promoter *P_tlpA_*, [34] all of which are compatible in both *E. coli* and *L. plantarum* WCFS1. The plasmid was constructed through Gibson assembly of the pLp-3050sNuc plasmid backbone and P*_tlpA_*-mCherry insert and transformed in *E. coli*. One correctly sequenced plasmid, extracted from an *E. coli* DH5α clone, was transformed in *L. plantarum* WCFS1 at different concentrations (300, 600, 900 and 1200 ng), yielding low TE values of 50 – 300 cfu/μg (Figure 2A) that increased with higher DNA concentrations. In the direct cloning method, the Gibson assembled plasmid was PCR amplified using complementary primers within the insert region and the resulting linear fragments were circularized by phosphorylation and ligation to yield sufficient quantities for transformation in *L. plantarum* WCFS1 (> 3μg). On the transformation of the circularized plasmid mix in *L. plantarum* WCFS1 at the different DNA concentrations as mentioned above, TE values were found to be lower than that of the indirect method although they were within the same order of magnitude and increased drastically when the DNA concentration was above 1 μg. The lower TE values could be due to incomplete circularization of the PCR-amplified plasmid fragments, due to which the final quantity of the circularized constructs might have been lower [35]. The accuracy of clones generated from the direct cloning method, determined by their ability to express mCherry was high at around 80% for all DNA quantities tested (Figure 2B). Notably, since *L. plantarum* WCFS1 contains 3 endogenous plasmids [36], sequencing-based verification of desired regions in the transformed plasmid was done by PCR amplifying the whole mCherry gene of 1140 base pairs (bp) (Figure 2C). The gene segments were directly amplified from bacterial cell pellets, and the amplicon sequencing was outsourced to an external provider, Eurofins Genomics GmbH, where their additional DNA purification option was employed. An initial purification of the PCR amplified product by us seemed to improve the quality of the sequencing chromatograms but was not absolutely necessary to get the correct results (Supporting information Figure S4A; Supporting information Figure S4B). As expected, all clones expressing mCherry yielded the correct sequences without any mutations or deletions.

**Figure 2.**
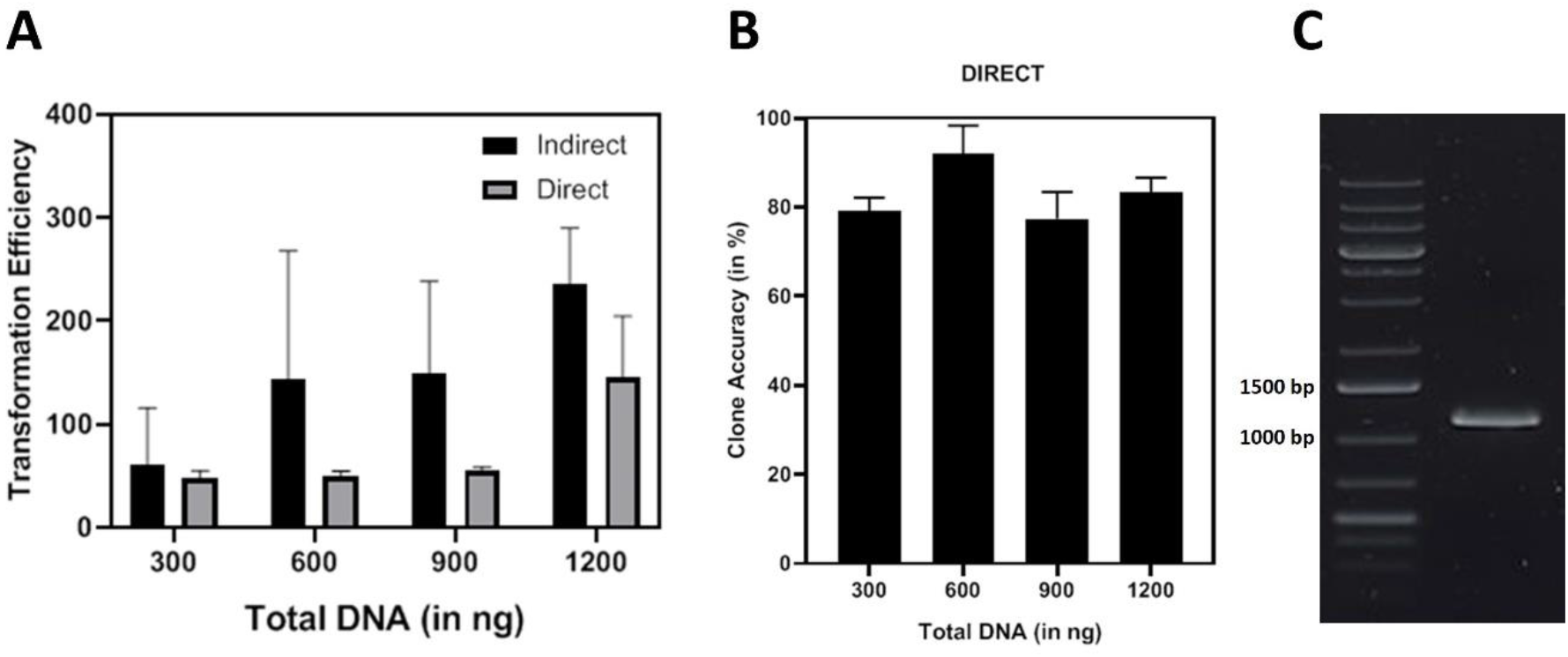
**A)** Transformation Efficiency comparison against the total concentration of DNA transformed for the indirect and Direct Mode of Cloning in *L. plantarum* WCFS1 (The standard deviations correspond to two independent biological replicates) **B)** Percentage accuracy of correct recombinant clones against the total concentration of DNA transformed in *L. plantarum* WCFS1 (The whiskers correspond to values from two independent biological replicates) C) Agarose gel showing the colony PCR product corresponding to the whole mCherry gene (1140 bp)

### Required Time for the direct and indirect cloning methods

The direct cloning method is considerably quicker and less labor-intensive than the indirect cloning method. All steps in the direct cloning method can be completed in 4 days after which PCR-amplified sequences can be sent for sequencing. In contrast, the indirect cloning method requires 5 to 6 days, depending on the time allocated for growth of bacteria on the master plate (Figure 3). Note that a master plate needed to be made from transformed *E. coli* colonies in our case due to the use of erythromycin as the common antibiotic resistance marker. *E. coli* has some natural resistance to this antibiotic because high doses of erythromycin are required to suppress the growth of non-transformed cells causing the transformed colonies to grow too small to be reliably handled for colony PCR. In the case of *L. plantarum* WCFS1, a master plate was not required since colonies grown for 48 hours were large enough to handle both colony PCR and culture inoculation.

**Figure 3.**
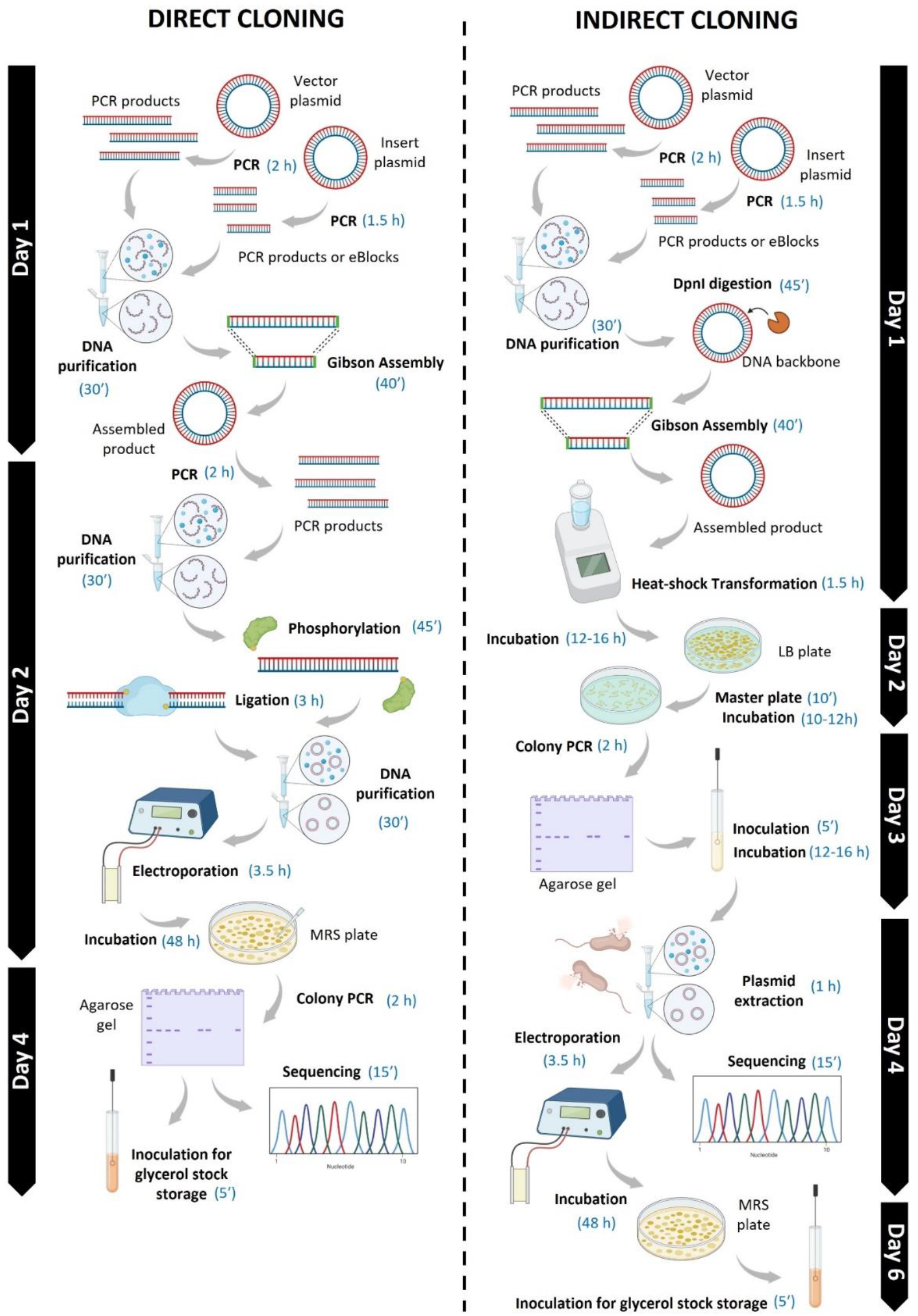
Schematic representation of the comparison of steps and time range between the direct and indirect cloning methods in *L. plantarum* WCFS1. The scheme was generated using BioRender.

### Direct cloning of a gene incompatible with *E. coli*

The main advantage of the direct cloning method is demonstrated in the ability to transform genes in lactobacilli that are challenging using the indirect method. Genes encoding proteins that are toxic to *E. coli*, for example, often result in mutations or complete deletions of the gene in the plasmid when transformed into *E. coli* [37]. We therefore tested the cloning of a plasmid containing the gene of a human peptidase inhibitor 3, elafin, encoded downstream of a strong constitutive promoter (*PtlpA*). This protease has been reported to exert anti-microbial activity with *E. coli*, so its constitutive expression is expected to be toxic [38,39]. Transformation of the assembled plasmid containing the constitutively expressed elafin in *E. coli* yielded very few colonies. The screening of positive clones was done by PCR amplification using a primer that annealed to the vector and one to the insert. Only 3 clones showed amplification, but the amplified product was shorter than expected (524 bp) (Figure 4A). Sequencing of plasmids extracted from these clones revealed several mutations and deletions. The whole *PtlpA* was deleted from all three plasmids, and two clones had the elafin coding sequence truncated (Figure 4B, Supporting information Figure S5). On the other hand, through the direct cloning method, over 124 colonies were obtained after transforming 1000 ng of phosphorylated and ligated product. 10 colonies were screened by PCR, and all showed amplification at the expected size of 524 bp(Figure 4A). Sequencing of the gene amplified from 3 clones revealed no mutations or deletions (Figure 4B).

**Figure 4.**
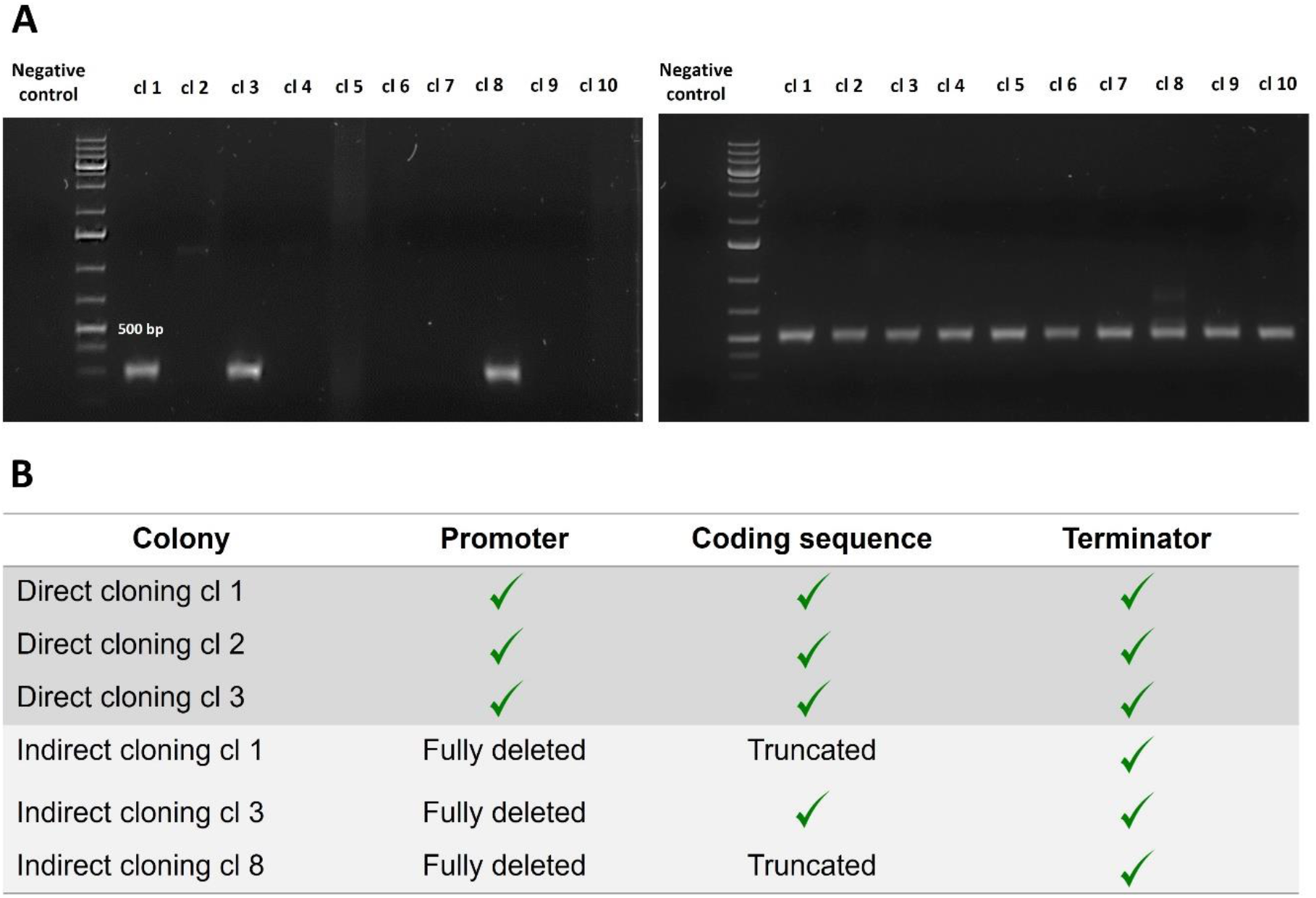
**A)** The agarose gel on the right shows the colony PCR result of 10 isolated *E. coli* colonies. Left agarose gel corresponds to the colony PCR of 10 isolated *L. plantarum* WCFS1 colonies. All colony PCRs were performed under the same conditions. The expected band size was 524 bp. The ladder used was the 1 kb Plus DNA ladder from ThermoFisher. **B)** Table listing the genetic mutations obtained in the elafin gene after the direct and indirect cloning methods (three clones per method)

While we have tested this direct cloning method only in *L. plantarum* WCFS1, we believe this strategy can also be applied to other hard-to-transform bacteria, providing that these bacteria accept unmethylated DNA.

## ACKNOWLEDGEMENTS

The plasmid *pLp3050sNuc* was a kind gift from Prof. Geir Mathiesen.

