## Supplementary tables and figures for "Gibson Assembly-based direct cloning of plasmid DNA in *Lactiplantibacillus plantarum* WCSF1"

| Primer Name | Sequence |
|---|---|
| **Vector mCherry fw** | 5’-GAAGATAAATCCCATAAGGG-3’ |
| **Vector mCherry rev** | 5’-AAGAACTCTATTGAAGCCC-3’ |
| **Insert mCherry fw** | 5’-GGGGCTTCAATAGAGTTCTTGGTGATGTCGGCGATATAG-3’ |
| **Insert mCherry rev** | 5’-CCCTTATGGGATTTATCTTCGACTCGCACTGAGAGGAT-3’ |
| **mCherry full amp fw** | 5’-TTACAAGGCTAAGAAGCCAG-3’ |
| **mCherry full amp rev** | 5’-GTAGTCTTAACTTCAGCATCG-3’ |
| **Vector elafin fw** | 5’-CATGGTATATCTCCTTCTT-3’ |
| **Vector elafin rev** | 5’-AATAACTAGCATAACCCC-3’ |
| **Insert elafin fw** | 5’-TAAGAAGGAGATATACCATGATGCGTGCTTCATCATTC-3’ |
| **Insert elafin rev** | 5’-AAGGGGTTATGCTAGTTATTTTAGTGATGGTGATGGTG-3’ |
| **Elafin full amp fw** | 5’-CAAGAAGTGTTGTGAAGGT-3’ |
| **Elafin full amp rev / Elafin colony PCR rev** | 5’-ATACCTGGACAATCAGTATCC-3’ |
| **Elafin colony PCR fw** | 5’-CAATGTTCCAAATGCGTG-3’ |
| **Whole gene sequencing fw** | 5’-AGTGGAACGAAAACTCAC-3’ |
| **Whole gene sequencing rev** | 5’-CGTTACTAAAGGGAATGGAG-3’ |

**Figure S1.** Primers used in this work.

**A**

GGGGCTTCAATAGAGTTCTTGGTGATGTCGGCGATATAGGCGCCAGCAACCGCACCTGTGGCGCCGGTGATGCCGGCCACGAT
GCGTCCGGCGTAGAGGATCGAGATCGATCTCGATCCCGCGAAATTTTAATTTGTTTGTTAGTTAGTTTATTTGTTGGTTTGTTTGT
GTTATAATATCCTCTAGAAATAATTTTGTTTAACTTTAAGAAGGAGATATACCATG**ATGGTTTCAAAGGGTGAAGAAGATAACATG**
**GCTATCATCAAGGAATTCATGCGTTTCAAGGTTCACATGGAAGGTTCAGTTAACGGTCACGAATTCGAAATCGAAGGTGAAG**
**GTGAAGGTCGTCCATACGAAGGTACTCAAACTGCTAAGTTAAAGGTTACTAAGGGTGGTCCATTACCATTCGCTTGGGATATCT**
**TATCACCACAATTCATGTACGGTTCAAAGGCTTACGTTAAGCACCCAGCTGATATCCCAGATTACTTAAAGTTATCATTCCCAGA**
**AGGTTTCAAGTGGGAACGTGTTATGAACTTCGAAGATGGTGGTGTTGTTACTGTTACTCAAGATTCATCATTACAAGATGGTG**
**AATTCATCTACAAGGTTAAGTTACGTGGTACTAACTTCCCATCAGATGGTCCAGTTATGCAAAAGAAGACTATGGGTTGGGAA**
**GCTTCATCAGAACGTATGTACCCAGAAGATGGTGCTTTAAAGGGTGAAATCAAGCAACGTTTAAAGTTAAAGGATGGTGGTC**
**ACTACGATGCTGAAGTTAAGACTACTTACAAGGCTAAGAAGCCAGTTCAATTACCAGGTGCTTACAACGTTAACATCAAGTTA**
**GATATCACTTCACACAACGAAGATTACACTATCGTTGAACAATACGAACGTGCTGAAGGTCGTCACTCAACTGGTGGTATGGAT**
**GAATTATACAAGTA**AAATAACTAGCATAACCCCTTGGGGCCTCTAAACGGGTCTTGAGGGGTTTTTTGCTGAAAGGAGGAACTA
TATCCGAACGATCCTCTCAGTGCGAGTCGAAGATAAATCCCATAAGGG

**B**

TATATAGGAGTATGATTCCC**ATGCGTGCTTCATCATTCTTAATCGTTGTTGTTTTCTTAATCGCTGGTACTTTAGTTTTAGAAGCT**
**GCTGTTACTGGTGTTCCAGTTAAGGGTCAAGATACTGTTAAGGGTCGTGTTCCATTCAACGGTCAAGATCCAGTTAAGGGTC**
**AAGTTTCAGTTAAGGGTCAAGATAAGGTTAAGGCTCAAGAACCAGTTAAGGGTCCAGTTTCAACTAAGCCAGGTTCATGTC**
**CAATCATCTTAATCCGTTGTGCTATGTTAAACCCACCAAACCGTTGTTTAAAGGATACTGATTGTCCAGGTATCAAGAAGTGTT**
**GTGAAGGTTCATGTGGTATGGCTTGTTTCGTTCCACAAGGTGGTTCACATCACCATCACCATCACTAA**TCTAGACTCGAGGAA
TTCGGTACCCCGGGTTCGAAGGCGCCAAGCTTCAAATTACAGCACGTGTTGCTTTGATTGATAGCCA

**Figure S2.** A) mCherry genetic sequence cloned in this work. The coding sequence is shown in bold. B) Elafin genetic sequence clones in this work.  sequence is shown in bold.

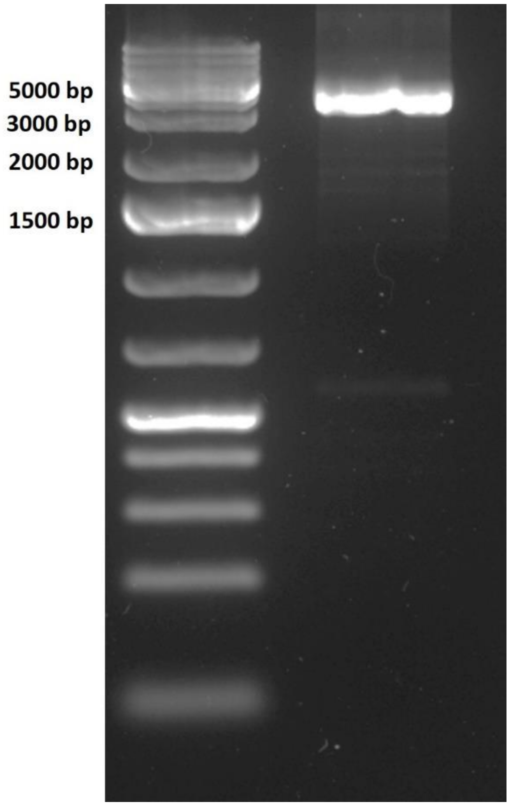


**Figure S3.** Agarose gel showing the HiFi PCR product (3728 bp). The 1kb Plus ladder was run as a reference (ThermoFisher).

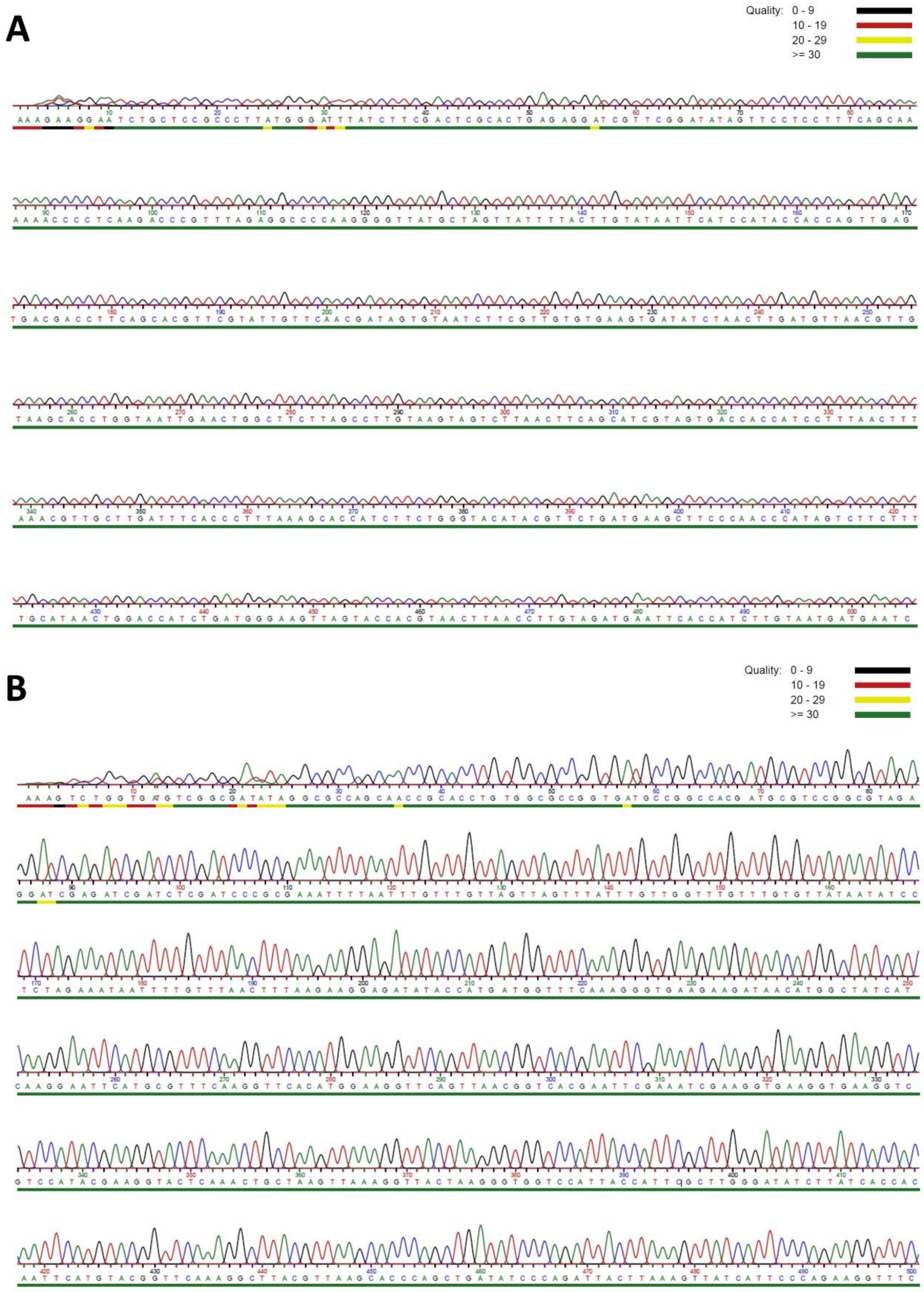


**Figure S4.** A) Sequencing chromatogram without an initial purification step. B) Sequencing chromatogram with an initial purification step.

$P_{TlpA}$ RBS Elafin CDS T7 ter

```
PTlpA_elafin        AGTAAGTTACAGGGCGAAGTAGTCAATGTTCCAAATGCGTGGTGATGTCGGCGATATAGG
E.coli_clone_1      AGTAAGTTACAGGGCGAAGTAGTCAATGTTCCAA--------------------------
                    **********************************

PTlpA_elafin        CGCCAGCAACCGCACCTGTGGCGCCGGTGATGCCGGCCACGATGCGTCCGGCGTAGAGGA
E.coli_clone_1      ------------------------------------------------------------

PTlpA_elafin        TCGAGATCGATCTCGATCCCGCGAAATTTTAATTTGTTTGTTAGTTAGTTTATTTGTTGG
E.coli_clone_1      ------------------------------------------------------------

PTlpA_elafin        TTTGTTTGTGTTATAATATCCTCTAGAAATAATTTTGTTTAACTTTAAGAAGGAGATATA
E.coli_clone_1      ------------------------------------------------------------

PTlpA_elafin        CCATGATGCGTGCTTCATCATTCTTAATCGTTGTTGTTTTCTTAATCGCTGGTACTTTAG
E.coli_clone_1      -----ATGCGTGCTTCATCATTCTTAATCGTTGTTGTTTTCTTAATCGCTGGTACTTTAG
                         *******************************************************

PTlpA_elafin        TTTTAGAAGCTGCTGTTACTGGTGTTCCAGTTAAGGGTCAAGATACTGTTAAGGGTCGTG
E.coli_clone_1      TTTTAGAAGCTGCTGTTACTGGTGTTCCAGTTAAGGGTCAAGATACTGTTAAGGGTCGTG
                    ************************************************************

PTlpA_elafin        TTCCATTCAACGGTCAAGATCCAGTTAAGGGTCAAGTTTCAGTTAAGGGTCAAGATAAGG
E.coli_clone_1      TTCCATTCAACGGTCAAGAT----------------------------------------
                    ********************

PTlpA_elafin        TTAAGGCTCAAGAACCAGTTAAGGGTCCAGTTTCAACTAAGCCAGGTTCATGTCCAATCA
E.coli_clone_1      --------------CCAGTTAAGGGTCCAGTTTCAACTAAGCCAGGTTCATGTCCAATCA
                                  **********************************************

PTlpA_elafin        TCTTAATCCGTTGTGCTATGTTAAACCCACCAAACCGTTGTTTAAAGGATACTGATTGTC
E.coli_clone_1      TCTTAATCCGTTGTGCTATGTTAAACCCACCAAACCGTTGTTTAAAGGATACTGATTGTC
                    ************************************************************

PTlpA_elafin        CAGGTATCAAGAAGTGTTGTGAAGGTTCATGTGGTATGGCTTGTTTCGTTCCACAAGGTG
E.coli_clone_1      CAGGTATCAAGAAGTGTTGTGAAGGTTCATGTGGTATGGCTTGTTTCGTTCCACAAGGTG
                    ************************************************************

PTlpA_elafin        GTTCACATCACCATCACCATCACTAAAATAACTAGCATAACCCCTTGGGGCCTCTAAACG
E.coli_clone_1      GTTCACATCACCATCACCATCACTAAAATAACTAGCATAACCCCTTGGGGCCTCTAAACG
                    ************************************************************

PTlpA_elafin        GGTCTTGAGGGGTTTTTTGCTGAAAGGAGG
E.coli_clone_1      GGTCTTGAGGGGTTTTTTGCTGAAAGGAGG
                    ******************************
```

```
PT1pA_elafin        AGTAAGTTACAGGGCGAAGTAGTCAATGTTCCAAATGCGTGGTGATGTCGGCGATATAGG
E.coli_clone_2      AGTAAGTTACAGGGCGAAGTAG-CAATGTTCCAA--------------------------
                    ********************** ***********

PT1pA_elafin        CGCCAGCAACCGCACCTGTGGCGCCGGTGATGCCGGCCACGATGCGTCCGGCGTAGAGGA
E.coli_clone_2      ------------------------------------------------------------

PT1pA_elafin        TCGAGATCGATCTCGATCCCGCGAAATTTTAATTTGTTTGTTAGTTAGTTTATTTGTTGG
E.coli_clone_2      ------------------------------------------------------------

PT1pA_elafin        TTTGTTTGTGTTATAATATCCTCTAGAAATAATTTTGTTTAACTTTAAGAAGGAGATATA
E.coli_clone_2      ------------------------------------------------------------

PT1pA_elafin        CCATGATGCGTGCTTCATCATTCTTAATCGTTGTTGTTTTCTTAATCGCTGGTACTTTAG
E.coli_clone_2      -----ATGCGTGCTTCATCATTCTTAATCGTTGTTGTTTTCTTAATCGCTGGTACTTTAG
                         *******************************************************

PT1pA_elafin        TTTTAGAAGCTGCTGTTACTGGTGTTCCAGTTAAGGGTCAAGATACTGTTAAGGGTCGTG
E.coli_clone_2      TTTTAGAAGCTGCTGTTACTGGTGTTCCAGTTAAGGGTCAAGATACTGTTAAGGGTCGTG
                    ************************************************************

PT1pA_elafin        TTCCATTCAACGGTCAAGATCCAGTTAAGGGTCAAGTTTCAGTTAAGGGTCAAGATAAGG
E.coli_clone_2      TTCCATTCAACGGTCAAGATCCAGTTAAGGGTCAAGTTTCAGTTAAGGGTCAAGATAAGG
                    ************************************************************

PT1pA_elafin        TTAAGGCTCAAGAACCAGTTAAGGGTCCAGTTTCAACTAAGCCAGGTTCATGTCCAATCA
E.coli_clone_2      TTAAGGCTCAAGAACCAGTTAAGGGTCCAGTTTCAACTAAGCCAGGTTCATGTCCAATCA
                    ************************************************************

PT1pA_elafin        TCTTAATCCGTTGTGCTATGTTAAACCCACCAAACCGTTGTTTAAAGGATACTGATTGTC
E.coli_clone_2      TCTTAATCCGTTGTGCTATGTTAAACCCACCAAACCGTTGTTTAAAGGATACTGATTGTC
                    ************************************************************

PT1pA_elafin        CAGGTATCAAGAAGTGTTGTGAAGGTTCATGTGGTATGGCTTGTTTCGTTCCACAAGGTG
E.coli_clone_2      CAGGTATCAAGAAGTGTTGTGAAGGTTCATGTGGTATGGCTTGTTTCGTTCCACAAGGTG
                    ************************************************************

PT1pA_elafin        GTTCACATCACCATCACCATCACTAAAATAACTAGCATAACCCCTTGGGGCCTCTAAACG
E.coli_clone_2      GTTCACATCACCATCACCATCACTAAAATAACTAGCATAACCCCTTGGGGCCTCTAAACG
                    ************************************************************

PT1pA_elafin        GGTCTTGAGGGGTTTTTTGCTGAAAGGAGG
E.coli_clone_2      GGTCTTGAGGGGTTTTTTGCTGAAAGGAGG
                    ******************************
```

```
PT1pA_elafin        AGTAAGTTACAGGGCGAAGTAGTCAATGTTCCAAATGCGTGGTGATGTCGGCGATATAGG
E.coli_clone_3      AGTAAGTTACAGGGCGAAGTAGT-------------------------------------
                    ***********************

PT1pA_elafin        CGCCAGCAACCGCACCTGTGGCGCCGGTGATGCCGGCCACGATGCGTCCGGCGTAGAGGA
E.coli_clone_3      ------------------------------------------------------------

PT1pA_elafin        TCGAGATCGATCTCGATCCCGCGAAATTTTAATTTGTTTGTTAGTTAGTTTATTTGTTGG
E.coli_clone_3      ------------------------------------------------------------

PT1pA_elafin        TTTGTTTGTGTTATAATATCCTCTAGAAATAATTTTGTTTAACTTTAAGAAGGAGATATA
E.coli_clone_3      ------------------------------------------------------------

PT1pA_elafin        CCATGATGCGTGCTTCATCATTCTTAATCGTTGTTGTTTTCTTAATCGCTGGTACTTTAG
E.coli_clone_3      ------------------------------------------------------------

PT1pA_elafin        TTTTAGAAGCTGCTGTTACTGGTGTTCCAGTTAAGGGTCAAGATACTGTTAAGGGTCGTG
E.coli_clone_3      -------------------------------------------TACTGTTAAGGGTCGTG
                                                               *****************

PT1pA_elafin        TTCCATTCAACGGTCAAGATCCAGTTAAGGGTCAAGTTTCAGTTAAGGGTCAAGATAAGG
E.coli_clone_3      TTCCATTCAACGGTCAAGATCCAGTTAAGGGTCAAGTTTCAGTTAAGGGTCAAGATAAGG
                    ************************************************************

PT1pA_elafin        TTAAGGCTCAAGAACCAGTTAAGGGTCCAGTTTCAACTAAGCCAGGTTCATGTCCAATCA
E.coli_clone_3      TTAAGGCTCAAGAACCAGTTAAGGGTCCAGTTTCAACTAAGCCAGGTTCATGTCCAATCA
                    ************************************************************

PT1pA_elafin        TCTTAATCCGTTGTGCTATGTTAAACCCACCAAACCGTTGTTTAAAGGATACTGATTGTC
E.coli_clone_3      TCTTAATCCGTTGTGCTATGTTAAACCCACCAAACCGTTGTTTAAAGGATACTGATTGTC
                    ************************************************************

PT1pA_elafin        CAGGTATCAAGAAGTGTTGTGAAGGTTCATGTGGTATGGCTTGTTTCGTTCCACAAGGTG
E.coli_clone_3      CAGGTATCAAGAAGTGTTGTGAAGGTTCATGTGGTATGGCTTGTTTCGTTCCACAAGGTG
                    ************************************************************

PT1pA_elafin        GTTCACATCACCATCACCATCACTAAAATAACTAGCATAACCCCTTGGGGCCTCTAAACG
E.coli_clone_3      GTTCACATCACCATCACCATCACTAAAATAACTAGCATAACCCCTTGGGGCCTCTAAACG
                    ************************************************************

PT1pA_elafin        GGTCTTGAGGGGTTTTTTGCTGAAAGGAGG
E.coli_clone_3      GGTCTTGAGGGGTTTTTTGCTGAAAGGAGG
                    ******************************
```

**Figure S5.** Multiple sequence alignment of the genetic sequence corresponding to the elafin gene, and the sequences obtained after Sanger sequences for each of the three isolated plasmids. All three clones have the full PTlpA and the RBS deleted. Clones 1 and 3 additionally have chunks of the elafin coding sequence deleted. The multiple sequence alignment tool MUSCLE was used to align the sequences.
